# Stratification of Chemotherapy-Treated Stage III Colorectal Cancer Patients Using Multiplexed Imaging and Single Cell Analysis of T Cell Populations

**DOI:** 10.1101/2021.02.24.432210

**Authors:** Xanthi Stachtea, Maurice B. Loughrey, Manuela Salvucci, Andreas U. Lindner, Sanghee Cho, Elizabeth McDonough, Anup Sood, John Graf, Alberto Santamaria-Pang, Alex Corwin, Pierre Laurent-Puig, Sonali Dasgupta, Sandra Van Schaeybroeck, Mark Lawler, Jochen H. M. Prehn, Fiona Ginty, Daniel B Longley

**Affiliations:** Patrick G. Johnston Centre for Cancer Research, School of Medicine, Dentistry and Biomedical Science, Queen’s University Belfast, Northern Ireland, UK; Department of Cellular Pathology, Royal Victoria Hospital, Belfast Health and Social Care trust, Belfast, UK; Department of Physiology and Medical Physics and Centre for Systems Medicine, Royal College of Surgeons in Ireland (RCSI) University of Medicine and Health Sciences, 123 St. Stephen’s Green, Dublin 2, Ireland; GE Research Center, 1 Research Circle, Niskayuna, NY, 12309, USA; UMR-S 1147, Université Paris Descartes, Paris, France; Velindre Cancer Centre, Velindre Rd, Cardiff, UK; Precision Medicine Centre of Excellence, The Patrick G Johnston Centre for Cancer Research, Queen’s University, Belfast, Northern Ireland, UK

**Author notes:** **Corresponding author**: Daniel B Longley, Patrick G. Johnston Centre for Cancer Research, School of Medicine, Dentistry and Biomedical Science, Queen’s University Belfast, Northern Ireland, UK.

## Abstract

Colorectal cancer (CRC) has one of the highest cancer incidences and mortality rates. In stage III, postoperative chemotherapy benefits <20% of patients, while more than 50% will develop distant metastases. Predictive biomarkers for identification of patients with increased risk for disease recurrence are currently lacking, with progress in biomarker discovery hindered by the disease’s inherent heterogeneity. The immune profile of colorectal tumors has previously been found to have prognostic value. The aims of this study were to evaluate immune signatures in the tumor microenvironment (TME) using an *in situ* multiplexed immunofluorescence imaging and single cell analysis technology (Cell DIVE™). Tissue microarrays (TMAs) with up to three 1mm diameter cores per patient were prepared from 117 stage III CRC patients treated with adjuvant fluoropyrimidine/oxaliplatin chemotherapy. Single sections underwent multilplexed immunofluorescence with Cy3- and Cy5-conjugated antibodies for immune cell markers (CD45, CD3, CD4, CD8, FOXP3, PD1) and cell segmentation markers (DAPI, pan-cytokeratin, AE1, NaKATPase and S6). We applied a probabilistic multi-class, multi-label classification algorithm based on multi-parametric models to build statistical models of protein expression to classify immune cells. Expert annotations of immune cell markers were made on a range of images, and Support Vector Machines (SVM) were used to derive a statistical model for cell classification. Images were also manually scored independently by a Pathologist as ‘high’, ‘moderate’ or ‘low’, for stromal and total immune cell content. Excellent agreement was found between manual and total automated scores (p<0.0001). Higher levels of multi-marker classified regulatory T cells (CD3+CD4+FOXP3+PD1-) were significantly associated with disease-free survival (DFS) and overall-survival (OS) (p=0.049 and 0.032), compared to FOXP3 alone. Our results also showed that PD1- Tregs rather than PD1+ Tregs were associated with improved survival. Overall, compared to single markers, multi-marker classification provided more accurate quantitation of immune cells with greater potential for predicting patient outcomes.

## Introduction

For early and locally advanced (stage I and II) colorectal cancer (CRC), the standard treatment of choice for low risk patients is surgical resection. Subsequent oncological treatment decisions for non-metastatic CRC are based largely on the anatomical AJCC/UICC TNM staging classification^1^. After the MOSAIC study in 2004, patients with stage III CRC now commonly receive oxaliplatin/fluoropyrimidine/leucovorin (5-fluorouracil (5FU), FOLFOX; or xeloda/capecitabine, XELOX) as standard adjuvant treatment ^2^. Of patients with stage III CRC treated with adjuvant chemotherapy, only ~20% will benefit from adjuvant FOLFOX, and 30% relapse within 2 to 3 years after surgery. Consequently, 80% of patients receive chemotherapy (and endure unnecessary toxicities) that yields no benefit ^3^. However, improvements in the understanding of CRC heterogeneity are paving the way for more personalized approaches that combine both histological and molecular data intelligence for patient stratification and therapy selection, including selecting which patients will benefit from adjuvant chemotherapy^4,5^.

In the past decade, there has been an increasing interest in the impact of the tumor microenvironment (TME) on patient prognosis. Decreased risk of tumor progression and improved survival have been observed in solid tumors with high T cell infiltration^6^. For CRC, the concept of an “Immunoscore” was introduced by Galon *et al;* this evaluates CD3- and CD8-positive immune infiltrates in the tumor core (TC) and tumor margin (TM) to classify “TNM-immune scores” for tumors^7^. In addition to Immunoscore, there have been numerous studies that reinforce the importance of tumor-infiltrating lymphocytes (TILs) as indicators of prognosis in CRC^8,9^. The importance of the immune contexture in CRC for patient prognosis logically suggests that immunotherapy could be a promising therapeutic approach^10^. Responsiveness to immunotherapy depends on several key factors, including high mutational loads (leading to high levels of tumor neoantigens), which are found in MMR-deficient (dMMR) microsatellite instability-high (MSI-high) CRC^11,12^. The immune checkpoint inhibitor (ICI) pembrolizumab has been approved by the US Food Drug Administration (FDA) for patients with metastatic dMMR/MSI-high CRC. However, the majority of colorectal tumors are microsatellite stable (MSS), with low mutational burdens and exhibit no response to ICI therapy. Thus, chemotherapy remains the backbone therapy for MSS CRC.

With the unmet clinical need to better stratify stage III patients for possible adjuvant (or neo-adjuvant) chemotherapy and the opportunity to better quantify immune response using newer cell quantification method, our goal was: 1) to compare multi-marker immune cell classification (using Cell DIVE) with immune cell scores determined by a pathologist and 2) investigate the association between single-marker and multi-marker immune cell classification and patient outcomes.

## Materials and Methods

### Patient Cohort

Tissue microarrays (TMAs) from formalin-fixed paraffin-embedded (FFPE) tissue blocks with up to three 1mm diameter cores per patient were prepared from 170 patients with stage III CRC. The punches were taken from the center of the tumor based on identification by a pathologist (Prof Manuel Salto-Tellez, Queen’s University Belfast). The patient samples were collected from three Research Centres: Beaumont Hospital (RCSI Hospital Group, Ireland), Queen’s University Belfast (UK) and Paris Descartes University (France) and the TMAs were constructed at Queen’s University Belfast. By design, the TMAs from 91 patients had 2 or 3 cores from each tumor. Pathological stage was determined by the AJCC TNM staging version applicable at the time of the reporting. All Centres provided ethical approval for this study and informed consent was obtained from all participants (NIB12-0034). Patients were recruited during 2005-2012.

At the patient level, the exclusion criteria based on tissue block or clinical data were: i) poor tissue quality or no tumor cells in tissue, ii) loss of follow-up or recurrence and/or death within less than two months from surgical resection, iii) absence of chemotherapy treatment, iv) positive resection margins, v) tumor site was appendix, vi) stage II or IV disease, vii) only one assessable core remaining after applying all exclusion criteria. At the tissue core level, individual cores on the TMA were excluded for assessment after pathology TMA slide review if no or minimal viable tumor was present for evaluation (i.e. minimal or no tumor tissue, heavily artefacted tissue, extensive tumor necrosis, extensive presence of normal adjacent tissue). After applying exclusion criteria from the original patient cohort, the remaining data comprised 117 stage III patients, who were all treated with 5FU-based adjuvant chemotherapy (predominantly FOLFOX or XELOX).

### *KRAS* status

A MassARRAY system (Sequenom) was used to detect somatic point mutations of *KRAS*.

### Multiplexed immunofluorescence analysis of TMAs

Multiplexed immunofluorescence staining of the CRC TMAs was performed as previously described^13^ using Cell DIVE™ Cytiva, Issaquah, WA), a multiplexed immunofluorescence microscopy method allowing for multiple protein markers to be imaged and quantified at cell level in a single tissue section. Briefly, formalin-fixed, paraffin-embedded (FFPE) tissue slides were de-paraffinized and rehydrated, underwent a two-step antigen retrieval and were then stained for 1 hour at room temperature using a Leica Bond autostainer. All antibodies were characterized per the previously described protocol^13^ and when possible, antibodies in routine clinical use were employed. After down-selection, each antibody was conjugated with either Cy3 or Cy5 bis-NHS-ester dyes using standard protocols as previously described^13^. All sections underwent multiplexed immunofluorescence for a total of 24 markers listed on **Supplementary Table 1**. The markers of interest for this study included CD3, CD4, CD8, FOXP3, CD45, NaKATPase, S6, pan-cytokeratin and AE1 and DAPI nuclear stain. All samples underwent DAPI imaging in every round, and background imaging for the first five rounds and every three rounds thereafter.

### Image processing, single cell segmentation

Using Cell DIVE automated image pre-processing software, all images were registered to baseline using DAPI and underwent autofluorescence subtraction, illumination and distortion correction. DAPI and Cy3 autofluorescence images were used to generate a pseudo-colored image, which visually resembles a Hematoxylin and Eosin (H&E) stained image, which we refer to as a virtual H&E (vH&E). This visualization format helps tissue QC review and facilitated review of tumor morphology and lymphocytes. All cells in the epithelial and stromal compartments were segmented using DAPI and pan-cytokeratin, while S6, and NaKATPase were used for subcellular analysis of epithelial cells. Each segmented cell was assigned an individual ID and spatial coordinate, as previously described^13^. Post segmentation, several quality control (QC) steps were conducted, including visual review and manual scoring of tissue quality and segmentation for every image, also described elsewhere^14^. Briefly, each image was reviewed for completeness and accuracy of segmentation masks in each subcellular compartment and tumor and stroma separation. Average biomarker intensity was calculated for each cell and the following additional cell filtering criteria were applied: 1) epithelial cells were required to have either 1-2 nuclei; 2) each sub-cellular compartment (nucleus, membrane, cytoplasm) area had to have > 10 pixels and < 1500 pixels; 3) cells had to have excellent alignment with the first round of staining (round 0); 4) cells were at >25 pixels distance from the image margins; 5) cell area for nuclear segmentation mask was >100 or <3000 pixels.

### Immune cell classification

A customized machine learning based algorithm^15^ developed as a Fiji (ImageJ) plug-in was used for immune cell classification. This is a probabilistic multi-class, multi-label classification algorithms based on multi-parametric models to build statistical models of protein expression and classify immune cells. The images were first segmented into epithelial and stromal regions or masks using a combination of PCK26 and AE1 (expressed in epithelial cells). Nuclei were segmented using DAPI signal and a wavelet-based algorithm^16^ and assigned to the epithelium or stromal regions based on co-localization of the nuclei with the epithelial or stromal masks. Manual expert annotations of the following markers associated with each segmented cell were made: AE1+, CD45+ CD3+, CD4+, CD8+, FOXP3+ and PD1+ and negative cells. Support Vector Machines (SVM) were used to derive a statistical model for cell classification. This multi-marker, annotation driven workflow was custom designed as an FIJI plug-in and allows analysis of complex multi-class models (up to 27 markers)^17,15^. Following classification, counts for both single-marker and multi-marker immune cell types were determined.

### Pathologist Scoring

A gastrointestinal pathologist (Maurice B. Loughrey, MBL) performed visual inspection of the virtual H&E slides generated from the DAPI and autofluorescence images^13,18^, for the 419 TMA cores. After applying exclusion criteria described earlier, 28 cores were excluded and 391 cores were assessed. MBL assigned two qualitative scores to each core comprising either ‘high’, ‘moderate’ or ‘low’ score, one for stromal cell content and one for immune cell content. For stroma, a high score was assigned when the stromal area was higher than the epithelial area; a moderate score was assigned when the stromal and the epithelial areas were equivalent; and a low score was assigned when the stromal area was lower than the epithelial area. The immune score was based on lymphoid cell abundance in the tissue core.

For equivalent comparison of the pathologist stroma and immune score with Cell DIVE automated scores the following steps were taken: 1) “Stromal cells” were defined as DAPI positive cells that were negative for all markers and outside the epithelial segmentation mask. The stroma score was calculated as the percentage of non-immune stromal cells in all segmented cells in the non-epithelial region; 2) “Immune cells” were defined as segmented cells that were positive for any of the immune markers (CD45, CD3, CD4, CD8) and negative for the AE1 epithelial marker. The immune scores were calculated from the counts of all segmented immune cells. 3) “Epithelial cells” were defined as segmented cells that were positive for AE1 staining and were within the Epithelial Segmentation Mask^19^.

### Statistical Analysis

For comparison of Cell DIVE quantitative stroma and immune scores with the pathologist scores, the Cell DIVE scores were categorized based on the pathologist’s three qualitative groups (high – moderate – low). Statistical analysis for comparison of group means was performed using Welch’s ANOVA and pairwise t-test. The association of the single-marker and multi-marker classified immune cells with clinical outcome was evaluated using both univariate and multivariate analyses with adjustment for clinico-pathological confounders (T, N, age, sex, nodal count, positive nodes, lymphovascular invasion, differentiation) in the multivariate Cox proportional hazards models. For the final multivariate models, the variables were subjected to backward elimination and the variables that did not contribute to model fit were removed. The final multivariate model was tested for multi-collinearity and proportional Hazards assumption. Variables with variance inflation factor > 2 were removed, and the remaining variables were re-subjected to backward elimination. The relative quality and goodness-of-fit of models was examined using Harrell’s C-index, and the model choice was determined by the Akaike Information Criterion (AIC). The T cell subtypes were counted and analysed as continuous variables after being transformed to ‘Percent of total’ tissue segmented cells, per patient. When the patients had multiple cores, the average percent of the assessable cores was calculated, unless stated otherwise. For survival analyses, the T cell subtypes calculated as % of total tissue cells were dichotomised at the median, and the Kaplan-Meier method was used to plot survival curves with the log-rank test used for comparisons. No adjustments were made for multiple comparisons. Hypothesis testing was performed at the 5% significance level.

The end-points studied were disease-free survival (DFS) and overall-survival (OS). DFS was the time between the study entry and either the date of the first recurrence, or the date that the last follow-up took place. OS was the time between the date of study entry and either the date of death from any cause, or the date of the last follow-up. All statistical analyses were performed in R Version 3.5.1 (https://cran.r-project.org).

## RESULTS

### Pathologist scoring versus automated immune cell classification

The TMA cores form the patients were assessed by the pathologist (MBL) and, after exclusion criteria, 62 patients had 3 assessable cores, 99 had 2 assessable cores whereas 7 patients had only 1 assessable core. Intra-tumor heterogeneity was reflected in intra-patient differences between the pathologist’s immune and stroma scores. Specifically, from the 62 patients with 3 assessable cores, only 13 (19%) had the same immune score and 18 (29%) the same stroma score for all three cores. For 5 (8%) patients, the immune score was different in each of the three cores, while for 6 (10%) patients, the stroma score was different in each of the three cores. This is to be expected given tumor histology variation in different core punches. From the 99 patients with two cores, 44 (44%) had the same immune score and 42 (42%) had the same stroma score in both tissue cores. In summary, for the 161 patients with more than one core, 104 (65%) showed immune heterogeneity and 101 (63%) showed stroma heterogeneity between their tissue cores. This highlights the inherent high degree of intra-tumor heterogeneity in CRC.

MBL performed visual inspection of the virtual H&E slides and assigned scores to each core of ‘high’, ‘moderate’ or ‘low’, for both stromal and immune cell content. We used the machine learning workflow to create a quantitative cell classification-based immune and stroma score (**Figure 1A**) to compare with the pathologist’s scores. The Cell DIVE immune (p < 0.001; **Figure 1B**) and stromal (p < 0.001; **Figure 1C**) score values were significantly associated with the corresponding pathologist’s scores. Therefore, the machine-learning-based Cell DIVE cell classification has potential to be used to evaluate tumor immune and stromal content.

**Figure 1:**
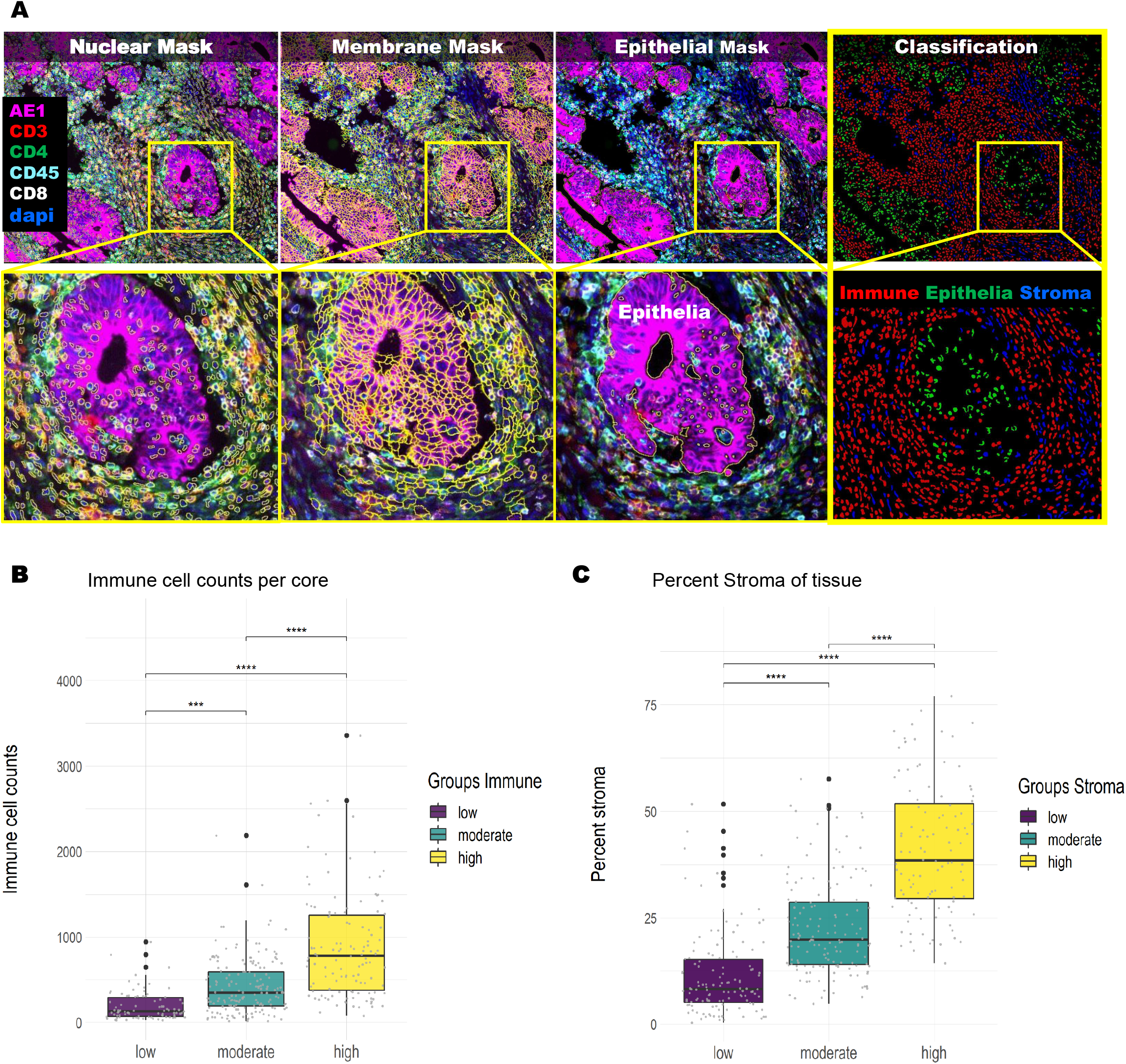
**A)** Cell DIVE workflow: immunofluorescence staining with overlaying segmentation masks and resulting classification. Based on the classification data, an immune and stroma score was calculated per TMA core. **B**.) The immune score was calculated from the immune cell density as counts of segmented cells that were positive for any of the immune markers (CD45, CD3, CD4, CD8) in each core. The cores were grouped based on the pathologist’s high-medium-low immune scores (x-axis) and each dot was the value of the Cell DIVE immune score (y-axis) per core. **C.)** The stroma score for each TMA core was calculated by counting segmented cells outside the epithelial mask that were negative for AE1 and immune markers, and converting to ‘percent of total’ cells. The cores were grouped based on the pathologist’s high-medium-low stroma scores (x-axis) and with each dot indicating stroma score (y-axis) per core. Statistical analysis was performed using Welch’s ANOVA and pairwise t-test (\*\*\**p* < 0.001 for all comparisons).

### T cell classification for single-marker and multi-marker (multiplexed) classification models

In order to study the impact of different T cell subtypes on patient prognosis in this adjuvant chemotherapy-treated cohort, we used a panel of T cell biomarkers as described earlier. In addition, to single marker analyses (CD3, CD4, CD8, FOXP3, PD1), multi-marker combinations were used to define subtypes (Tc, TcPD1, Th, ThPD1, Treg, TregPD1, **Figure 2A**). In the single-marker classification workflow, each one of these immune markers was analysed individually, and each segmented cell was classified as either positive or negative for each marker. Since the individual markers are used to generate the multi-marker classification, it is not surprising that they were significantly correlated (p<0.001; **Supplementary Figure 3**). The demographic data of the patient cohort are summarized in **Table 1**.

**Figure 2:**
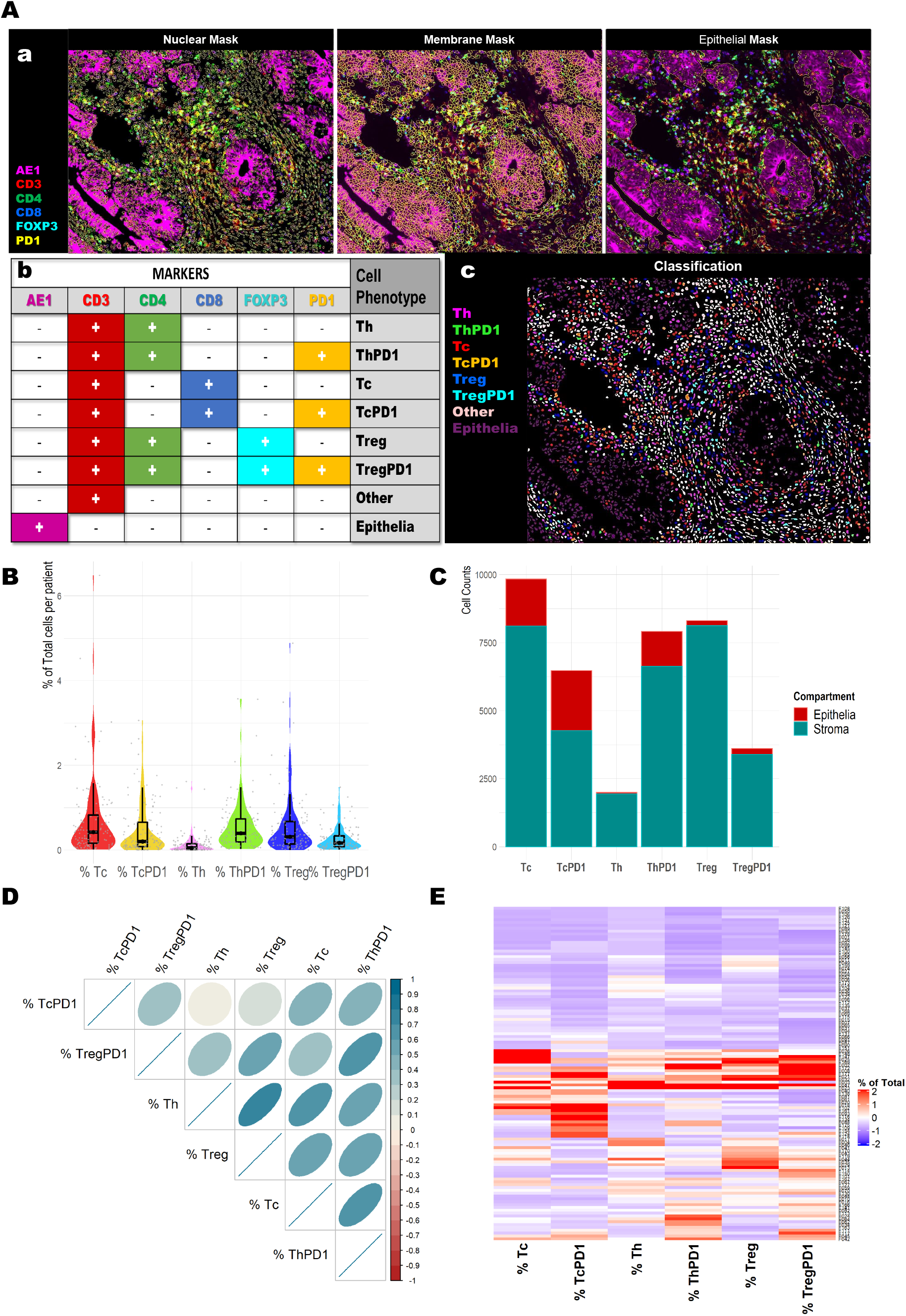
**A(a)** Representative multiplexed immunofluorescent tissue images and segmentation masks that were used for the multi marker classification workflow. For single marker classification, the immune markers were assessed individually (CD3, CD4, CD8, FOXP3, PD1) and each cell was classified as positive or negative for each marker. **A(b)** Marker combination (AE1, CD3, CD4, CD8, FOXP3, PD1) was used for multi-marker classification workflow and based on the marker co-staining each cell was assigned a cell subtype as shown in the table. The eight multi marker T cell subtypes were: T helper cells (**Th**), T helper PD1 (**ThPD1**), T cytotoxic (**Tc**), T cytotoxic PD1+ (**TcPD1**), T regulatory (**Treg**), T regulatory PD1 positive (**TregPD1**), **Epithelial** and **Other** (non-lymphocyte, non-epithelial) cells**. A(c)** Illustration of the resulting multi-marker classification for the nuclear mask. **B)** Distribution of calculated values for % of Total for each T cell subtype, per patient. Each value represents the average of 2-3 assessable cores, per patient. **C)** Total counts from all patients grouped as epithelial-associated cells (red) and stromal cells (green) for each multi-marker T cell subtype. **D)** Correlation matrix showing the relationship between different T cell subtypes (Spearman’s correlation coefficients). A color-coded correlation scale is provided and blue ellipses represent positive correlations, while darker color and narrower ellipses correspond to larger correlation coefficient magnitudes. E) Heatmap showing separation and clustering of patients based on % T cells of Total cells in tumor cores. Clusters based on the Ward.D agglomerative clustering method with Euclidean correlation distance measure.

**Table 1:**
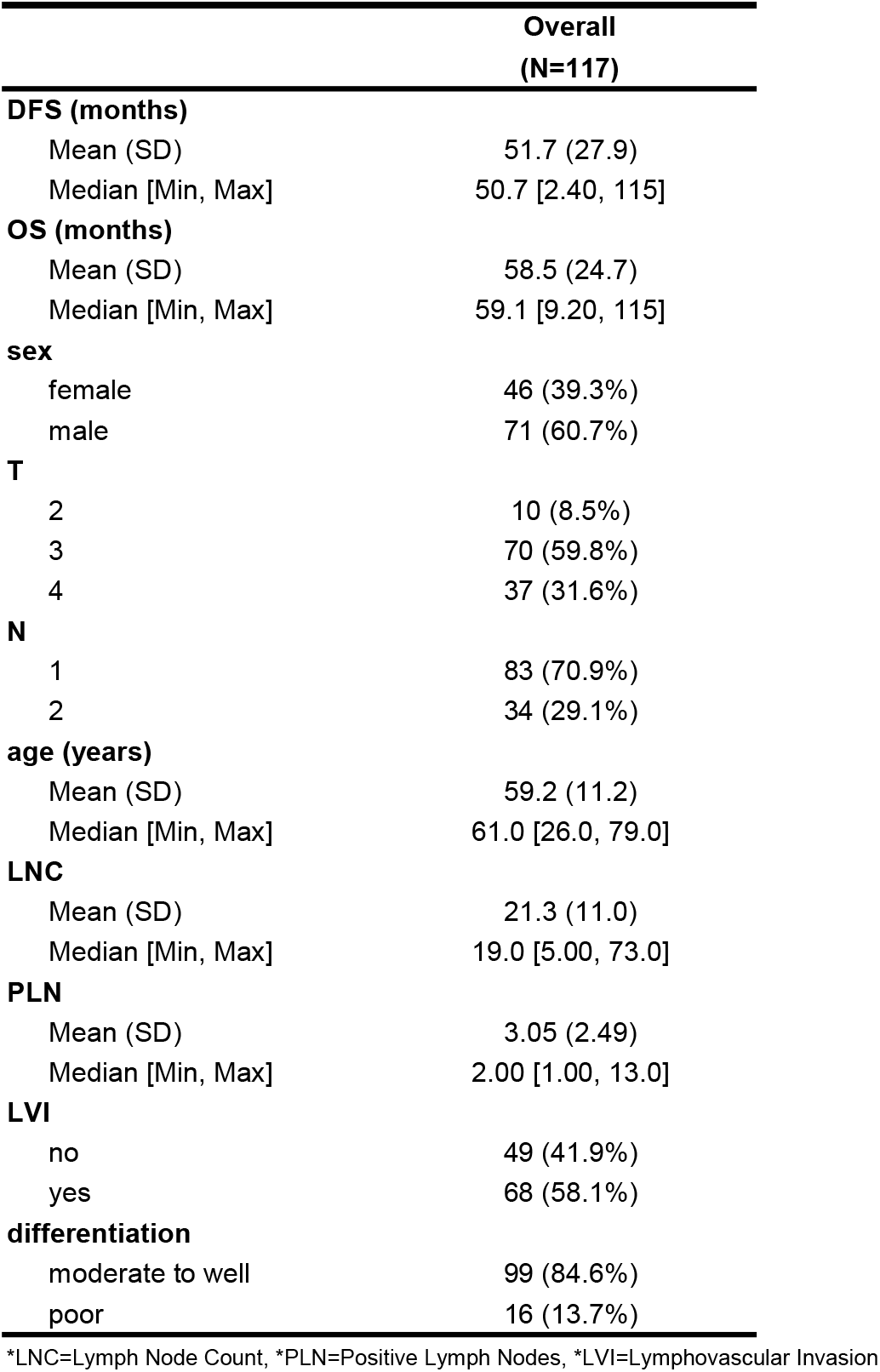
Demographic data of patient cohort

Representative immunofluorescent images of a single tissue core for the individual markers and the corresponding Segmentation Masks are illustrated in **Supplementary Figure 1**. In the multi-marker classification workflow all markers were assessed simultaneously (**Figure 2A(a)**) and, depending on marker co-localization, segmented cells were assigned to the following classes (**Figure 2A(b)**): PD1-negative T-helper (Th), PD1-positive Th (ThPD1), PD1-negative cytotoxic T cells (Tc), PD1-positive Tc (TcPD1), PD1-negative Treg and PD1-positive Treg (TregPD1).

To account for tumor heterogeneity, only patients with more than 1 core were used for the analysis (117 patients). Each T cell subtype was calculated as a percentage of total cells per core and the average percentage per patient was calculated. The distribution of T cell subtypes across the cohort is shown in **Figure 2B**; Tc and TcPD1 cells were the most abundant subtype associated with the epithelial compartment; however, overall and as expected, the majority of each T cell subtype was located in the stroma (**Figure 2C**). All T cell subtypes were generally positively correlated with each other, except TcPD1 had minimal correlation with Th and Treg (**Figure 2D**). Hierarchical clustering was used to assess the immune landscape of the patient cohort (**Figure 2E**). Separation into two clusters, immune “hot” (higher immune cells) and “cold” (lower immune cells), showed that nearly 50% of patients were low in all T cell subtypes; however, Kaplan-Meier analyses showed that their prognosis was similar to patients with higher level of T cells (**Supplementary Figure 2A**).

After separating into three clusters, the “immune-hot” cluster of patients with highest infiltration of T cell subtypes showed improved DFS and OS compared to the other 2 groups that had lower T cell levels; however this did not reach statistical significance (**Supplementary Figure 2B**). Detailed summary statistics for T cells for the multi-marker classifications and single marker classifications are presented in **Table 2**.

**Table 2:** Summary statistics for multi-marker and single-marker subtypes.

|  | Overall<br>(N=117) |
| --- | --- |
| **% Tc (of total cells)** |  |
| Mean (SD) | 0.693 (0.925) |
| Median [Min, Max] | 0.423 [0.0111, 6.49] |
| **% TcPD1** |  |
| Mean (SD) | 0.477 (0.619) |
| Median [Min, Max] | 0.197 [0, 3.06] |
| **% Th** |  |
| Mean (SD) | 0.133 (0.251) |
| Median [Min, Max] | 0.0511 [0, 1.62] |
| **% ThPD1** |  |
| Mean (SD) | 0.559 (0.593) |
| Median [Min, Max] | 0.394 [0.0222, 3.57] |
| **% Treg** |  |
| Mean (SD) | 0.568 (0.732) |
| Median [Min, Max] | 0.312 [0, 4.88] |
| **% TregPD1** |  |
| Mean (SD) | 0.262 (0.275) |
| Median [Min, Max] | 0.166 [0.00471, 1.48] |
| **% CD3** |  |
| Mean (SD) | 3.18 (2.88) |
| Median [Min, Max] | 2.52 [0.221, 22.3] |
| **% CD4** |  |
| Mean (SD) | 3.64 (3.52) |
| Median [Min, Max] | 2.67 [0.126, 23.3] |
| **% CD8** |  |
| Mean (SD) | 2.17 (2.42) |
| Median [Min, Max] | 1.35 [0.0283, 12.3] |
| **% FOXP3** |  |
| Mean (SD) | 1.58 (1.46) |
| Median [Min, Max] | 1.19 [0.0114, 8.57] |
| **% PD1** |  |
| Mean (SD) | 0.669 (0.694) |
| Median [Min, Max] | 0.395 [0.00943, 3.41] |

In **Supplementary Figure 4** representative images of virtual H&Es, immunofluorescent images and tissue mappings with color-coded cell classifications are illustrated. The selected images are representative of all 9 Stroma-Score/Immune-Score combinations from the pathologist review. This shows that multiplexing can be used to identify multiple subtypes of immune cells simultaneously, allowing for associations and potential cross-talk between distinct cell subtypes in the TME to be assessed.

### T cell infiltration and patient prognosis

As proof-of-concept for the applicability of this approach for identification of prognostic immune biomarkers, we next determined the prognostic value of the single and multiplexed markers in this FOLFOX-treated stage III patient cohort. The correlation of each T cell type with clinical endpoints (DFS and OS) was analysed using univariate and multivariate Cox proportional hazards models and Kaplan-Meier analyses.

In the univariate analyses, the forest plots in **Figure 3** demonstrate that none of the single immune markers was significantly associated with DFS (**Figure 3A**) or OS (**Figure 3B**), whereas the level of Treg cells (CD3+/CD4+/FOXP3+/PD1-) from the multi-marker machine-learning classification was significantly associated with longer DFS *(HR = 0.37, 95% CI = 0.14-0.99, p = 0.047)*.

**Figure 3:**
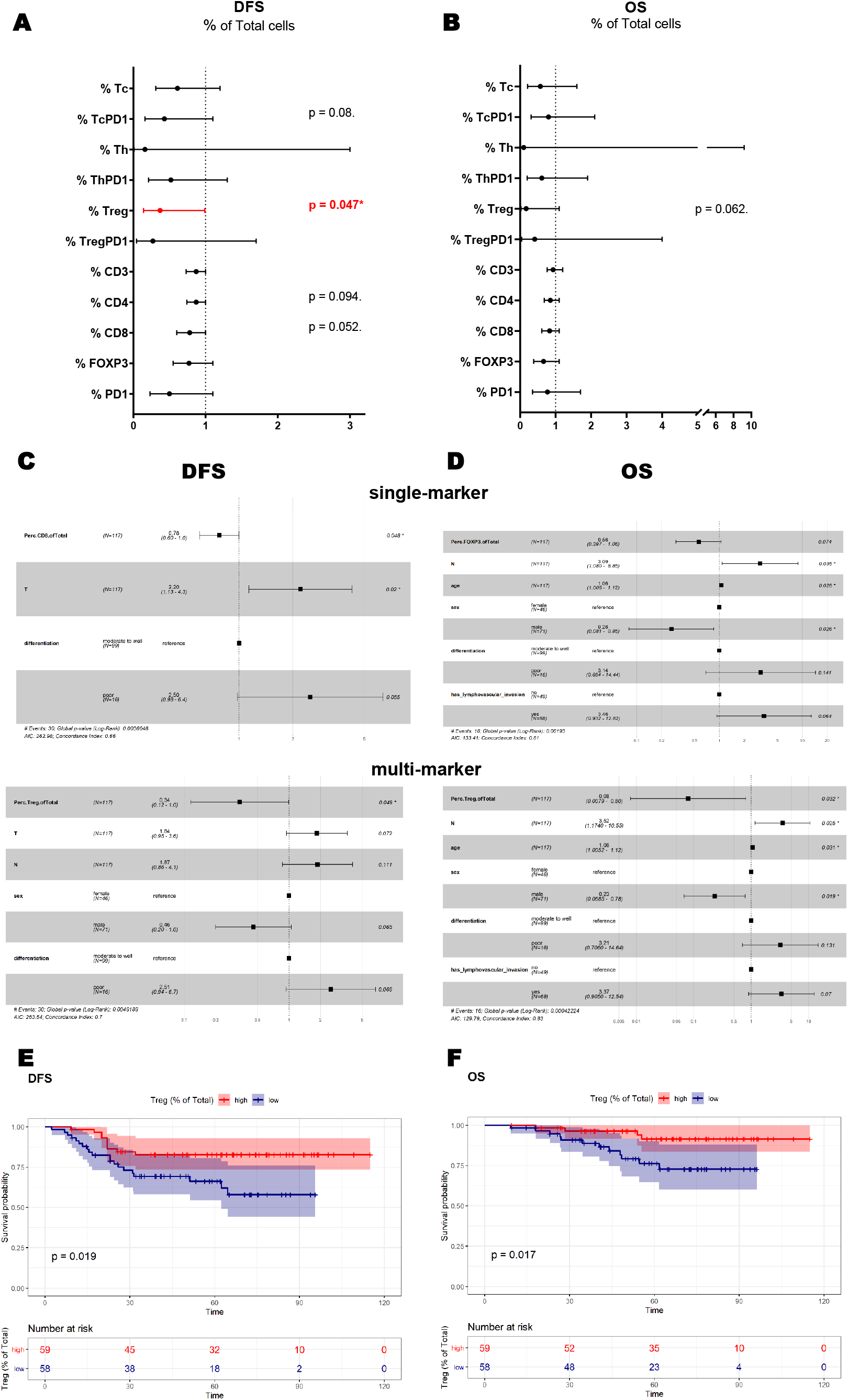
Survival estimates for DFS and OS for **Average** of cores. Forest plots for multi-marker classification (Tc, TcPD1, Th, ThPD1, Treg, TregPD1) and single-marker classification (CD3, CD4, CD8, FOXP3, PD1) for the patient cohort. Estimated HRs, 95% CIs, and p values from likelihood ratio tests from univariate Cox proportional hazards models demonstrated the associations between the percent of total classified cells with the risks of recurrence (DFS) (**A**) **and** death (OS) (**B**). In the multivariate analysis the biomarkers were adjusted for clinical variables (T, N, age, sex, nodal count, positive nodes, differentiation, lymphovascular invasion) for DFS (**C**) and OS (**D**). Kaplan–Meier curves demonstrating univariate survival analysis for percent of total for Treg dichotomized at the median for DFS (**E**) and OS (**F**). Differences in Kaplan–Meier survival curves are presented as log- rank P value.

For the multivariate analysis, the model initially included the clinical variables: T, N, age, sex, nodal count, positive nodes, differentiation and lymphovascular invasion together with single- and multi-marker immune scores. Backward elimination was used to select variables for the final model. For DFS in the single-marker model, CD8 remained in the final model and was positively associated with longer DFS (*multivariate adjusted HR = 0.78, 95% CI = 0.6 - 1.0, p = 0.048;* **Figure 3C**) and in the multi-marker model Tregs remained positively associated with longer DFS (*multivariate adjusted HR = 0.34, 95% CI = 0.12 - 1.0, p = 0.049;* **Figure 3C**). For OS in the single-marker model, FOXP3 remained in the final model but did not reach significance (*multivariate adjusted HR = 0.56, 95% CI = 0.297 – 1.06, p = 0.074;* **Figure 3D**) and in the multi-marker model Tregs remained positively associated with longer OS (*multivariate adjusted HR = 0.08, 95% CI = 0.0079 – 0.8, p = 0.032;* **Figure 3D**). The detailed Forest plots for the multivariate models for clinical variables only are shown in **Supplementary Figure 5.**

In order to facilitate comparison with previously published results, Treg levels were divided into high and low groups using the sample median as the cut-off and Kaplan-Meier analyses were performed for curves for DFS and OS (**Figure 3E** and **F**). Similar to the univariate and multivariate analyses above, *Treg-high* patients had improved DFS (*p = 0.019*) and OS (*p = 0.017*) than Treg*-low* patients. Kaplan-Meier curves for all single-marker and multi-marker classes dichotomized on the median are included in **Supplementary Figure 6**. Sub-regional analysis based on the percentage of immune cell subtypes located in the stroma or located within/associated with the epithelial compartment and association with outcome are shown in **Supplementary Table 2**.

### T cell infiltration and patient prognosis for immune hot-spot

In order to account for tumor immune heterogeneity, the average percent T cells in multiple cores was used for the above data analyses. However, this could dilute the impact of very high but very localised immune cell infiltrates. We hypothesised that by focusing our analyses on the available cores with *highest* tumor immune regions, we might uncover additional prognostic information; therefore, we repeated the above analyses for the one core per patient with maximum T cell density for each subtype.

We calculated the total counts of T cells in each core (CD3 counts for single markers and sum of all T cell subtypes for the multiplexed model). From the 117 patients, the cores with the highest number of CD3 or T cells (immune hot-spot core) was selected for further analysis. Cox proportional hazards regression analysis and Kaplan-Meier plots were performed as above. In the univariate analysis none of the single markers was significantly associated with survival. For the multi-marker classification Treg levels were significantly associated with DFS (*HR = 0.51, 95% CI = 0.27-0.97, p = 0.04;* **Figure 4A**) and were borderline significant for OS (*HR = 0.24, 95% CI = 0.059-1, p = 0.05;* **Figure 4B**).

**Figure 4:**
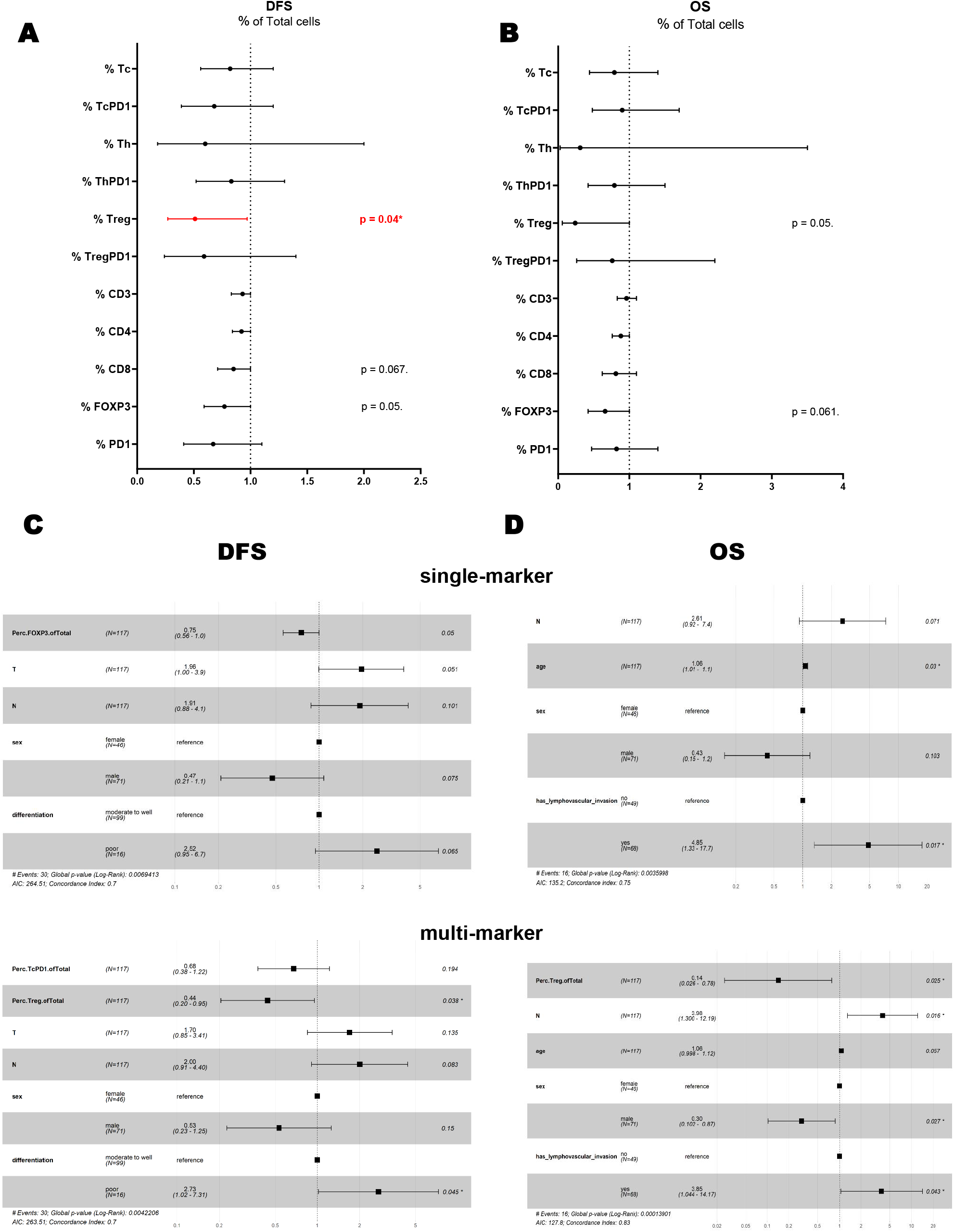
Survival estimates for DFS and OS for **immune hot-spot**: Forest plots for multi-marker classification (Tc, TcPD1, Th, ThPD1, Treg, TregPD1) and single-marker classification (CD3, CD4, CD8, FOXP3, PD1) for the patient cohort. Estimated HRs, 95% CIs, and P values from likelihood ratio tests from univariate Cox proportional hazards models demonstrated the associations between the percent of total classified cells with the risks of recurrence (DFS) (**A**) and death (OS) (**B**). In the multivariate analysis the biomarkers were adjusted for clinical variables (T, N, age, sex, nodal count, positive nodes, differentiation, lymphovascular invasion) for DFS (**C**) and OS (**D**).

In the multivariate analysis, for DFS in the single-marker model, FOXP3 remained in the final model (*multivariate adjusted HR = 0.75, 95% CI = 0.56-1.0, p = 0.05*) and had borderline statistical significance (**Figure 4C**), and in the multi-marker model Treg and TcPD1 remained in the final model and Treg remained statistically significant (for TcPD1: *multivariate adjusted HR = 0.68, 95% CI = 0.38-1.22, p = 0.194;* for Treg: *multivariate adjusted HR = 0.44, 95% CI = 0.20-0.95, p = 0.038*). For OS, none of the single markers remained in the final model. In the multi-marker model, Treg levels remained in the final model and were significantly associated with improved OS *(multivariate adjusted HR = 0.14, 95% CI = 0.026-0.78, p = 0.025*) (**Figure 4D**).

As previously, Kaplan-Meier curves for all single marker and multi-marker classes dichotomized on the median are included in **Supplementary Figure 7.** Sub-regional analysis based on the percentage of immune cell subtypes located in the stroma or located within/associated with the epithelial compartment and association with outcome are shown in **Supplementary Table 3**.

### Association of *KRAS* status with survival and distribution of T cell subtypes

In adjuvant FOLFOX/XELOX-treated stage III colorectal cancer patients, *KRAS* mutations have been associated with shorter time to recurrence (TTR) and OS^20^. We performed survival analysis for the patients with known *KRAS* status (108 out of 117 patients tested) to study the effect of this mutation in our cohort and its interaction with T cell subtype levels. Survival curves for colorectal cancer DFS and OS were plotted using the Kaplan-Meier method and compared by the log rank test using the *KRAS* mutation status as the stratification variable. *KRAS* status was not significantly associated with prognosis in our cohort, although, interestingly, there was a non-significant trend (p = 0.07) for *KRAS* mutant tumors to be associated with better DFS in this FOLFOX-treated cohort (**Supplementary Figure 8**). *KRAS* mutation has also been reported to have an immunosuppressive effect in the tumor microenvironment of colorectal cancer^21^. Summary statistics for the clinicopathological data of the patients grouped by *KRAS* status are shown in **Table 3**. No differences of T cell subtypes were observed between *KRAS* WT and mutant patients, for any of the classes tested for multiplexed classification or single marker classification. Collectively, these results indicate that the prognostic impact of T cell subtypes is not associated with *KRAS* mutational status.

**Table 3:** Demographic data and summary statistics of multi-marker model based on KRAS status

|  | WT<br>(N=64) | mutant<br>(N=44) | p-value |
| --- | --- | --- | --- |
| **sex** |  |  |  |
| female | 24 (37.5%) | 20 (45.5%) | 0.53 |
| male | 40 (62.5%) | 24 (54.5%) |  |
| **T** |  |  |  |
| 2 | 4 (6.2%) | 6 (13.6%) | 0.342 |
| 3 | 37 (57.8%) | 26 (59.1%) |  |
| 4 | 23 (35.9%) | 12 (27.3%) |  |
| **N** |  |  |  |
| 1 | 42 (65.6%) | 34 (77.3%) | 0.277 |
| 2 | 22 (34.4%) | 10 (22.7%) |  |
| **age scores** |  |  |  |
| <50 | 18 (28.1%) | 7 (15.9%) | 0.296 |
| >70 | 14 (21.9%) | 7 (15.9%) |  |
| 51-60 | 11 (17.2%) | 11 (25.0%) |  |
| 61-70 | 21 (32.8%) | 19 (43.2%) |  |
| **LNC** |  |  |  |
| <12 | 8 (12.5%) | 5 (11.4%) | 0.919 |
| >20 | 28 (43.8%) | 21 (47.7%) |  |
| Dec-20 | 28 (43.8%) | 18 (40.9%) |  |
| **PLN** |  |  |  |
| N>7 | 6 (9.4%) | 2 (4.5%) | 0.393 |
| N1-3 | 42 (65.6%) | 34 (77.3%) |  |
| N4-6 | 16 (25.0%) | 8 (18.2%) |  |
| **LVI** |  |  |  |
| no | 21 (32.8%) | 23 (52.3%) | 0.0683 |
| yes | 43 (67.2%) | 21 (47.7%) |  |
| **% Tc (of total cells)** |  |  |  |
| Mean (SD) | 0.636 (0.728) | 0.848 (1.20) | 0.355 |
| Median [Min, Max] | 0.421 [0.0111, 3.02] | 0.475 [0.0199, 6.49] |  |
| **% TcPD1** |  |  |  |
| Mean (SD) | 0.550 (0.670) | 0.411 (0.546) | 0.606 |
| Median [Min, Max] | 0.215 [0, 3.06] | 0.254 [0, 2.63] |  |
| **% Th** |  |  |  |
| Mean (SD) | 0.107 (0.215) | 0.191 (0.309) | 0.0721 |
| Median [Min, Max] | 0.0393 [0, 1.62] | 0.115 [0, 1.47] |  |
| **% ThPD1** |  |  |  |
| Mean (SD) | 0.547 (0.538) | 0.634 (0.695) | 0.63 |
| Median [Min, Max] | 0.407 [0.0222, 3.56] | 0.416 [0.0564, 3.57] |  |
| **% Treg** |  |  |  |
| Mean (SD) | 0.525 (0.669) | 0.680 (0.862) | 0.371 |
| Median [Min, Max] | 0.303 [0.0319, 3.49] | 0.352 [0.00996, 4.88] |  |
| **% TregPD1** |  |  |  |
| Mean (SD) | 0.287 (0.312) | 0.245 (0.227) | 0.778 |
| Median [Min, Max] | 0.182 [0.00471, 1.48] | 0.166 [0.0253, 0.910] |  |

\*LNC=Lymph Node Count, \*PLN=Positive Lymph Nodes, \*LVI=Lymphovascular Invasion

## DISCUSSION

Currently 5FU-based chemotherapy (usually FOLFOX or XELOX) is used as adjuvant treatment for stage-III CRC patients^2^. However, only 20% of patients benefit, while 30% will experience recurrence^3^. Therefore, reliable biomarkers that can predict which stage III patients would benefit from adjuvant chemotherapy is an urgent unmet clinical need in CRC.

A large number of multigene signatures using tumor gene expression profiles have emerged in the last decade, such as Consensus Molecular Subgroups (CMS) and CRC Intrinsic Subtypes (CRIS), which classify patients into molecular subtypes for risk prediction^22,23,24^. However, this approach is therapeutically valuable only under the assumption that highest risk patients will also be the most responsive to chemotherapy. This is not the case and, in fact, CMS4 patients who are predicted to have poor prognosis do not benefit from intensive adjuvant chemotherapy^25^. We recently reported that stage-II patients with CMS2/CRIS-C tumors, which demonstrate low levels of CD8-positive tumor-infiltrating lymphocytes benefit from adjuvant chemotherapy. In stage III patients, benefit from chemotherapy was particularly apparent in CMS2/CRIS-C and CMS2/CRIS-D patients^5^. However, transcriptional profiling is not routinely available or applied in clinical practice. Ideally, a clinical test to triage patients for adjuvant chemotherapy that could be performed rapidly on a single FFPE tumor section would be extremely useful.

Herein, we explored the potential of the Cell DIVE platform for enumerating several key immune cell populations previously linked with patient outcome in CRC. Using FFPE tissue samples, Cell DIVE can measure up to 60 markers within a single histological section, whereas standard IHC would require multiple sections to achieve similar result. This requirement would introduce the problem of cellularity changes through the sequential sections, as well as require extensive use of often limited valuable biological material. In addition, with Cell DIVE, multiple markers can be visualized simultaneously, thus increasing specificity for cell classification and providing a molecular signature within a histological content. In concert with user-friendly machine-learning methodologies, Cell DIVE has the potential to become a routine digital pathology platform for clinical laboratory settings. To demonstrate this, we compared immune and stroma scoring from the visual inspection of all tissues by a gastrointestinal pathologist with the corresponding Cell DIVE-derived immune and stroma scoring. Cell DIVE scoring showed significant association with the pathologist’s scores, suggesting that the Cell DIVE platform provides robust immune and stroma scoring for tumor tissue sections.

Using the Cell DIVE platform and a segmentation and classification workflow involving 10 markers, we show that we were able to detect 6 classes of T cells and associate them with patient prognosis using a single section from TMAs of FOLFOX/XELOX-treated stage III patients. Our results showed that high levels of Treg cells (CD3+/CD4+/FOXP3+/PD1-) were associated with improved survival in this cohort and were distinct from their PD1+ counterparts.

Treg cells are key mediators of self-tolerance, regulating multiple immune cells, such as CD4+ and CD8+ effector cells, macrophages and dendritic cells^26^. In the thymus, CD25+/CD4+ thymocytes can become Treg precursors, which, after stimulation with IL-2 and TGF-β, will differentiate into natural thymic FOXP3+ Tregs^27,28^. Natural Tregs can recognize self-antigens and migrate to damaged tissues to supress the activity of other T cells and prevent an uncontrolled inflammatory response^29,30^. Outside the thymus, in secondary lymphoid organs and peripheral tissues, Tregs are derived from differentiation of naïve conventional CD4+ T cells in response to cytokines that induce FOXP3 expression^31,32^. In CRC, there are higher levels of Tregs in the tumor than in healthy tissue. Recently, it has been shown that tumor-associated Tregs have distinct differences from normal peripheral Tregs^33,34^. In cancer, Tregs can suppress anti-tumor immune responses^35^ or have protective roles by controlling cancer-associated inflammation^36,37^. Within the intestine, immune cells reside within the mucosa^38^ and are tightly associated with the intestinal microbiome, thus intestinal Treg depletion can lead to unresolved inflammation^28,36^.

High Treg levels have been associated with poor clinical outcomes in different cancers, including CRC^35,39,40^, in contrast to our findings; other studies have associated high Treg levels with better prognosis in CRC patients^41,42,43,44,45^. There are a number of reasons that could be responsible for these apparently contradictory results. For example, differences in the study cohorts, most notably, stage and whether patients were treated with chemotherapy, but also cohort size, variable thresholds for scoring, technical differences in detection and scoring between laboratories and different follow-up times^46^ may contribute to these findings. In addition, the conflicting results may be due to lack of robust biomarkers that can reflect the Treg versatility and plasticity, and the best classification method for Treg is still under active debate. FOXP3 is a biomarker with high selectivity for Treg identification that is routinely used as a Treg biomarker in clinical studies. However, it has limitations since it is not exclusively expressed by Treg cells. FOXP3 can also be expressed in dividing, activated T effector cells^47,48^. Apart from FOXP3, some Treg subtypes can express other molecules that increase their immuno-suppressive capacity, and these highly suppressive Treg cells have been detected in CRC patients^49,50,51^. Therefore, relying solely on FOXP3 as a marker of Tregs may be the cause of some of the inconsistencies in the literature regarding Treg and CRC prognosis.

Recently, it was shown that the majority of intra-tumoral T cells in the TME are CD4+ with co-expression of PD1 molecules^52,53^, which is similar to our findings. PD-1 expression on T cells can be sign of early activation or exhaustion and reduced effector functions, due to prolonged exposure to tumor antigens^54^. Our results show that PD1- Tregs rather than PD1+ Tregs are associated with improved survival. Overall, the presence of Treg correlates with the presence of T effector cells in inflamed tissues. Given that CRC is a highly inflammatory type of tumor, PD1-Treg enrichment may not be associated with pro-tumorigenic immunosuppression, but rather are recruited as a result of an active immune response; this would explain the association which we observed with improved prognosis in this chemotherapy-treated stage-III cohort. Lastly, one limitation of our study is the lack of untreated patients for comparison. Therefore, we cannot evaluate the potential of Tregs as a prognostic biomarker. Of note, Tregs have been associated with better prognosis both for treated^41^ and untreated patients^55^.

The association of T cell infiltration with patient prognosis was assessed using: i) the average of two or three available TMA cores, and ii) the core with the highest T cell infiltration (or the “immune hot-spot” core). While using the core average can better account for TME heterogeneity and may be more representative of an entire tumor section, the immune hot spot core could better reveal subtle immune signatures of distinct T cell subtypes and infiltration levels that would have otherwise been attenuated or lost by averaging. Comparing the two workflows, the results were similar, especially in the univariate analysis, where none of the single markers was significant, while Treg cells were significantly associated with DFS in both workflows. In the multivariate analysis, the results were comparable for the multi-marker classes, with Treg cells remaining significant, while the single markers were inconsistent between workflows. In addition, when using hierarchical clustering for the immune hot-spot core we were unable to discover a distinct immune signature that was significantly associated with survival (data not shown). Since the TMA cores were randomly selected from the tumor center, it is unknown whether the immune hot -pot represents the entire tumor or is random event. Considering these limitations, averaging may be a superior sampling approach for assessing tumor immune infiltration than selecting the hot-spot for the purposes of this study.

*KRAS* status has been reported to be a biomarker for outcome in MSS stage III CRC^20^. *KRAS* mutational status was not significantly associated with outcome in our study, although there was a non-significant association for improved DFS in the *KRAS* mutant group. *KRAS* mutation has been associated with an immunosuppressive TME in MSS CRC^21^. However, no differences were observed in the levels of any the T cell subtypes examined between *KRAS* WT and mutant tumors in our study.

In summary, we show that multiplexed analysis of tissue and multi-marker cell classification can be used to accurately determine immune cells in tumor and stroma in colorectal tumor cores. We also provide proof-of-concept evidence for its utility to identify highly specific immune subsets that have clinical relevance.

Supplementary information is available at *Modern Pathology’s* website

## Acknowledgements

Research reported in this publication was partially supported by the National Cancer Institute of the National Institutes of Health under award number R01CA208179 supporting FG, EMcD, AS, JG and AS-P and AC. DBL and XS were supported by a US-Ireland R01 award (NI Partner supported by HSCNI, STL/5715/15). JHMP is supported by Science Foundation Ireland and the Health Research Board (16/US/3301). ML is supported by Health Data Research UK

## Conflict of Interest

ML has received honoraria from Pfizer, EMD Serono and Roche for presentations unrelated to this work. ML is supported by an unrestricted educational grant from Pfizer for research unrelated to this work.

---

**Supplementary Figure 1:** Panel of representative multiplexed image with the corresponding segmentation masks and individual staining for each antibody used for T cell classification and segmentation in this study.

**Supplementary Figure 2:** Heatmap showing separation and clustering of patients based on % T cells of Total T cell subtypes in tumor cores. Clusters based on the Ward.D agglomerative clustering method with Euclidean correlation distance measure. Kaplan-Meier curves color-coded for the corresponding 2 (A) and 3 (B) clusters of patients, demonstrating univariate survival analysis for DFS and OS. Differences in Kaplan–Meier survival curves are presented as log- rank p value.

**Supplementary Figure 3:** Pearson Correlation between cell counts for single marker model and multi marker classification. **CD3**: CD3 vs Tc+TcPD1+Th+ThPD1+Treg+TregPD1, **CD4**: CD4 vs Th+ThPD1+Treg+TregPD1, **CD8**: CD8 vs Tc+TcPD1, **FOXP3**: FOXP3 vs Treg+TregPD1, **PD1**: PD1 vs TcPD1+ThPD1+TregPD1. The average counts of % of Total per patient were used for the analysis.

**Supplementary Figure 4:** Representative vHE images and the corresponding multiplexed immunofluorescence and tissue mappings after classification.

**Supplementary Figure 5:** Forest plots of multivariate Cox proportional hazards models for clinical variables after backward elimination.

**Supplementary Figure 6:** Univariate survival analysis using Kaplan-Meier curves for single-marker and multi-marker T cell subtypes as percent of Total for average. The patients were separated in two groups using the median as cut-off.

**Supplementary Figure 7:** Univariate survival analysis using Kaplan-Meier curves for single-marker and multi-marker T cell subtypes as percent of Total for immune hot-spot cores. The patients were separated in two groups using the median as cut-off.

**Supplementary Figure 8: A)** Association of Kras status with survival for treated patients for DFS and OS.

