## SupplementalMaterial for "Stratification of Chemotherapy-Treated Stage III Colorectal Cancer Patients Using Multiplexed Imaging and Single Cell Analysis of T Cell Populations"

Supplementary Figure 1

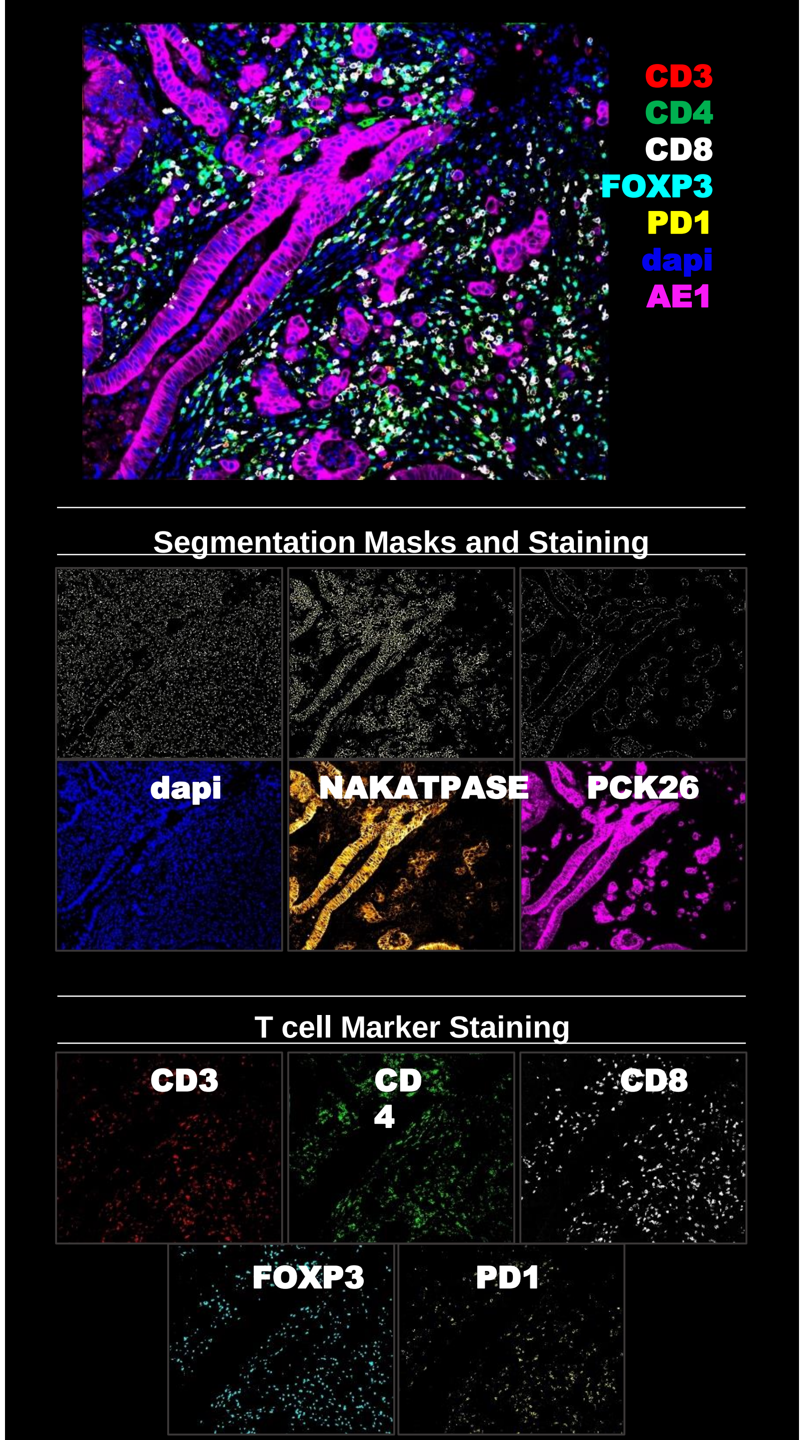

Supplementary Figure 2

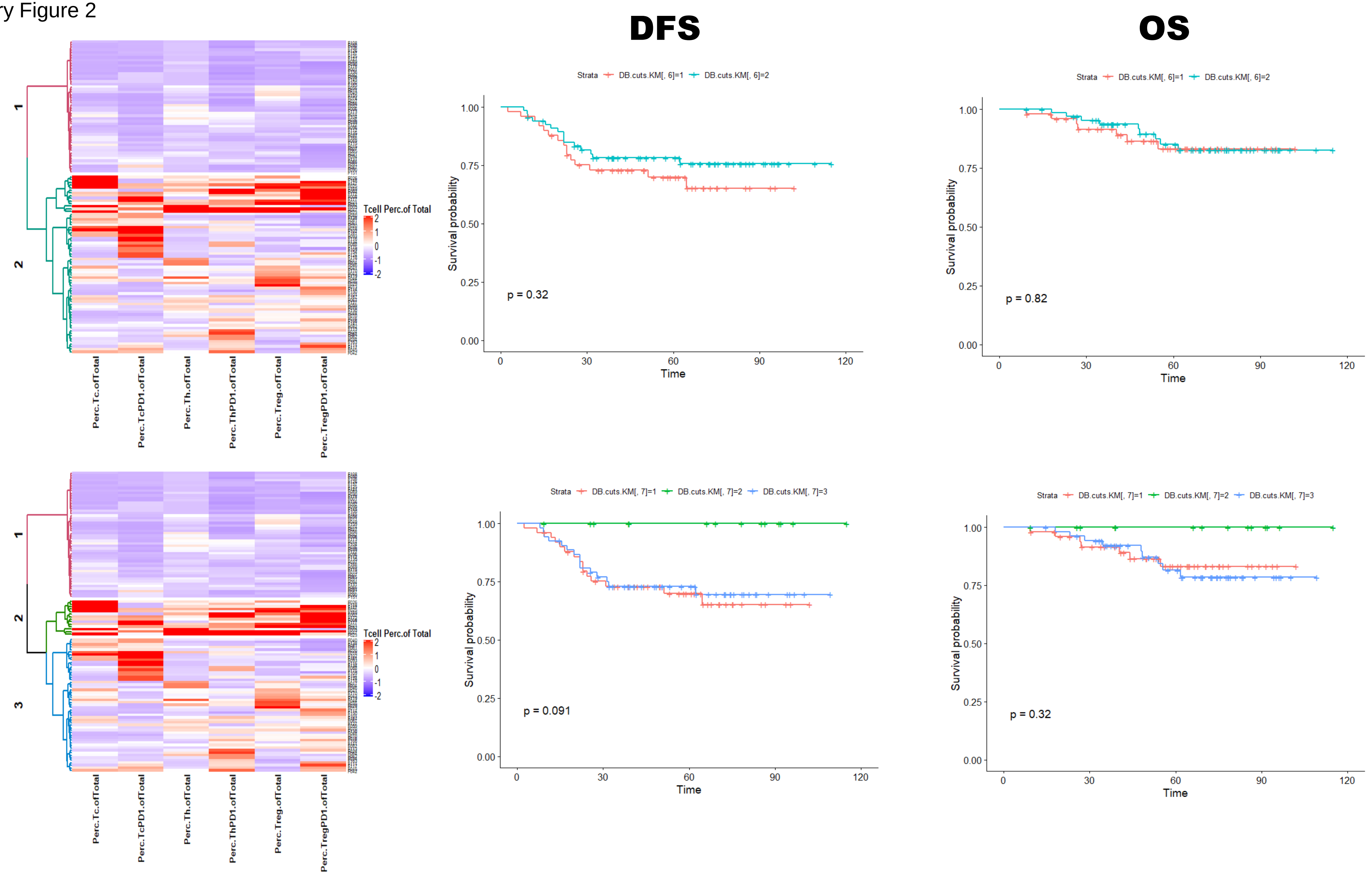

Supplementary Figure 3

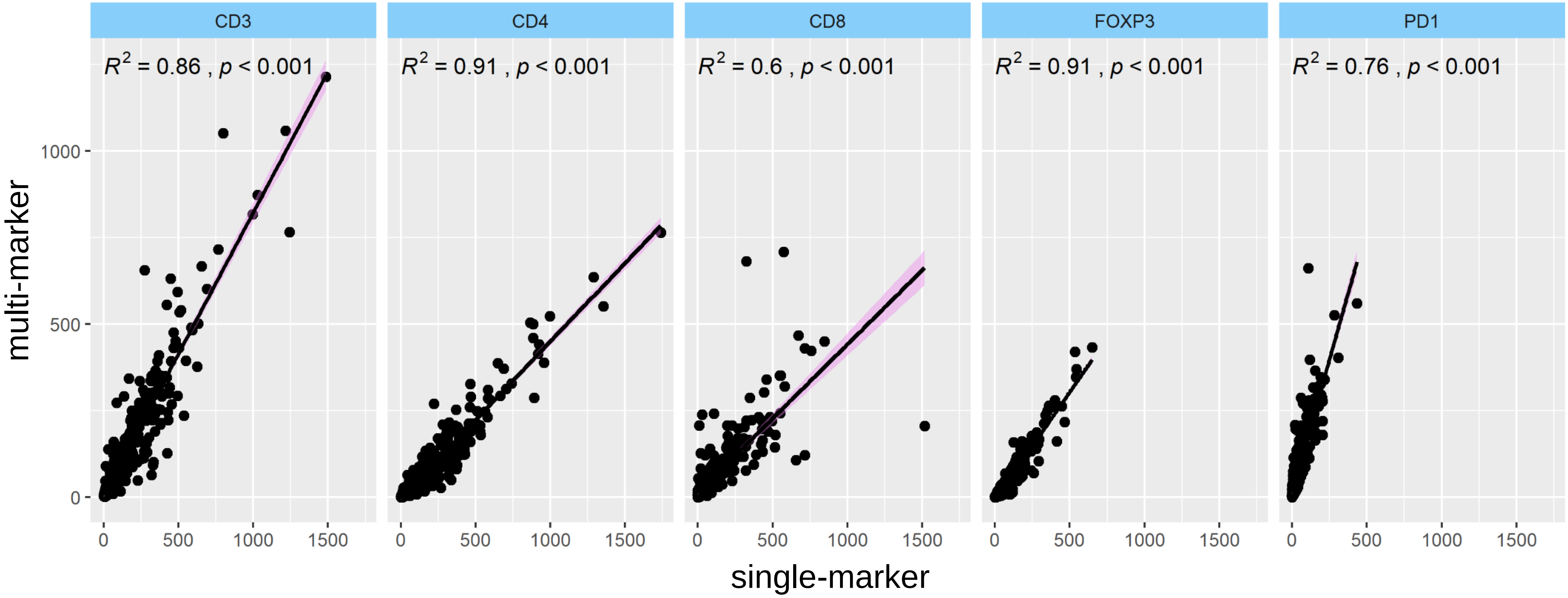

Supplementary Figure 4

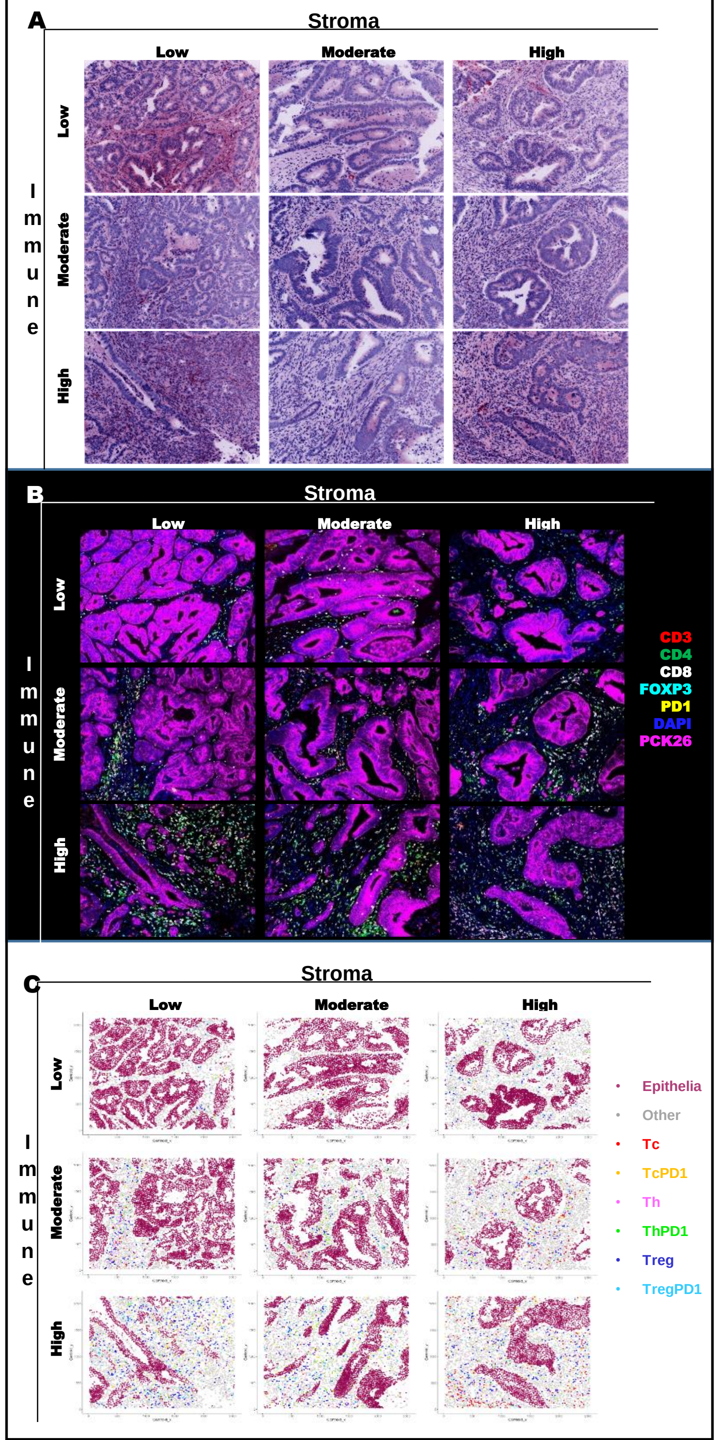

Supplementary Figure 5

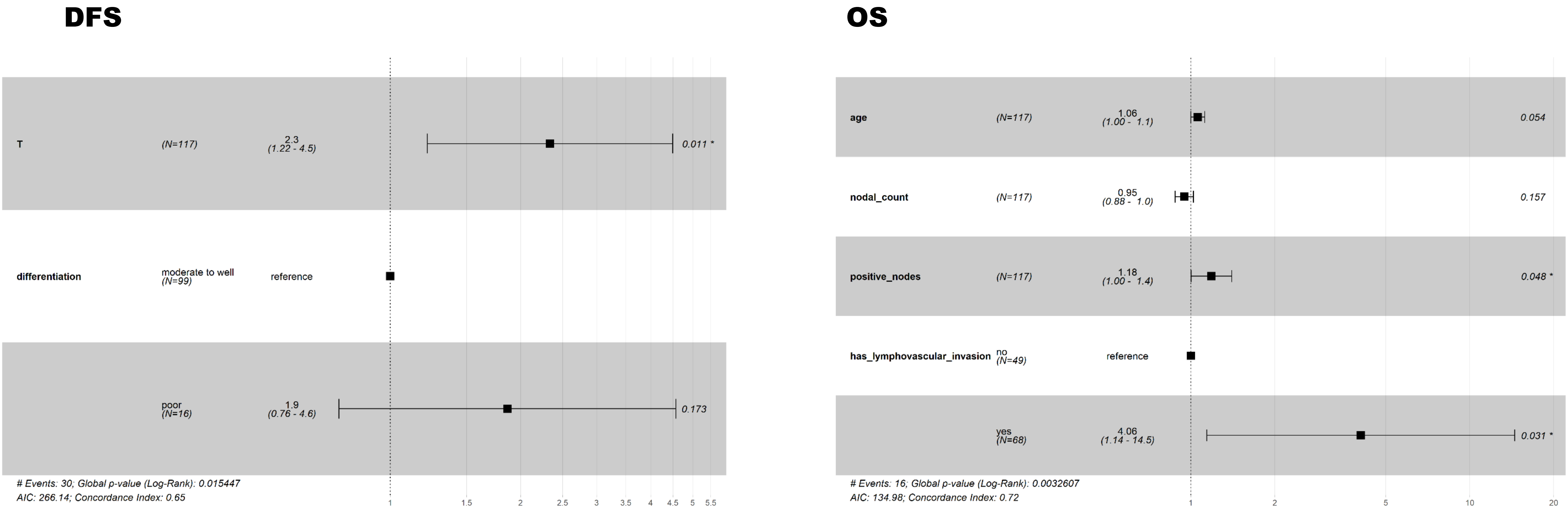

Supplementary Figure 6

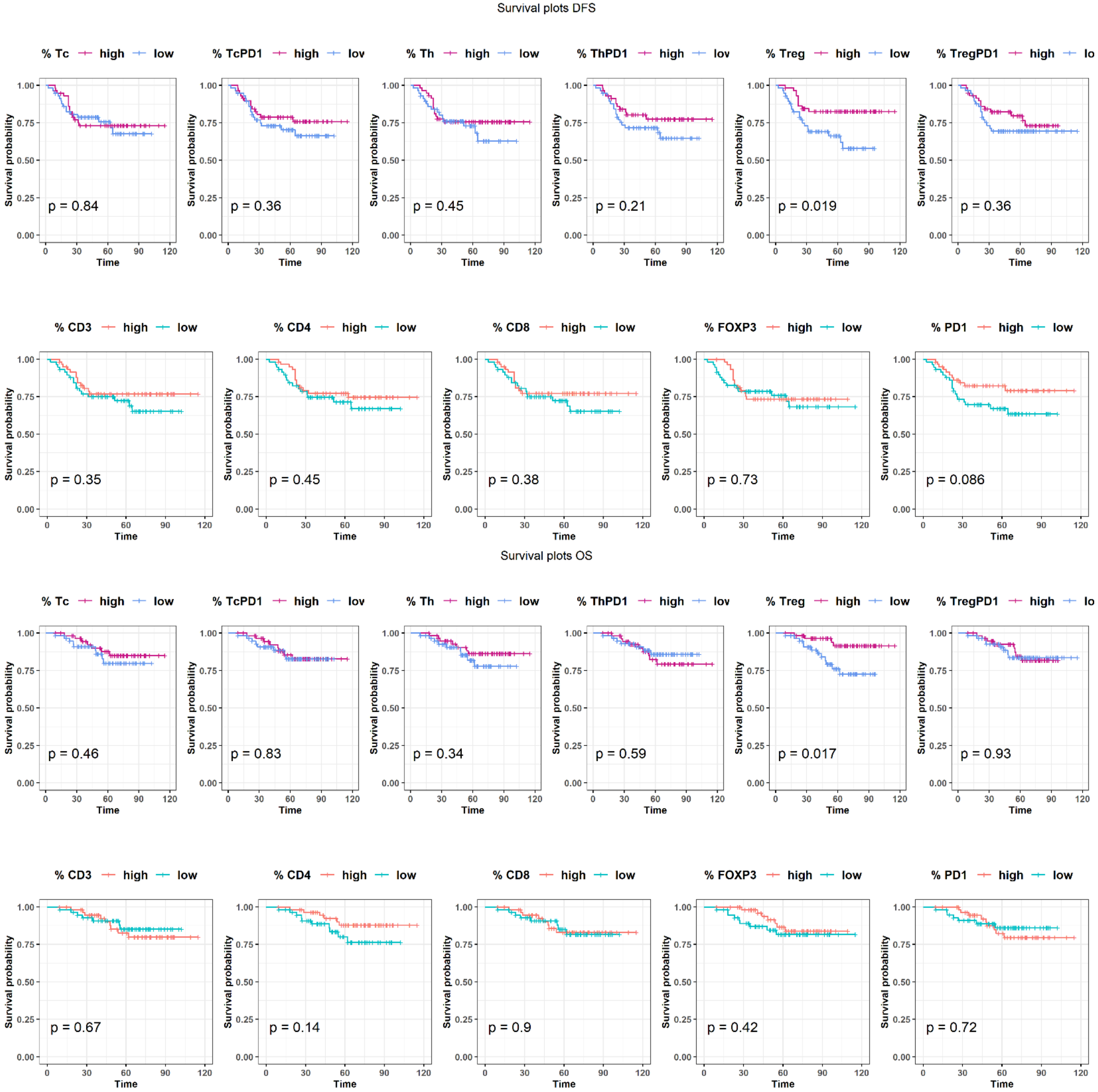

Supplementary Figure 7

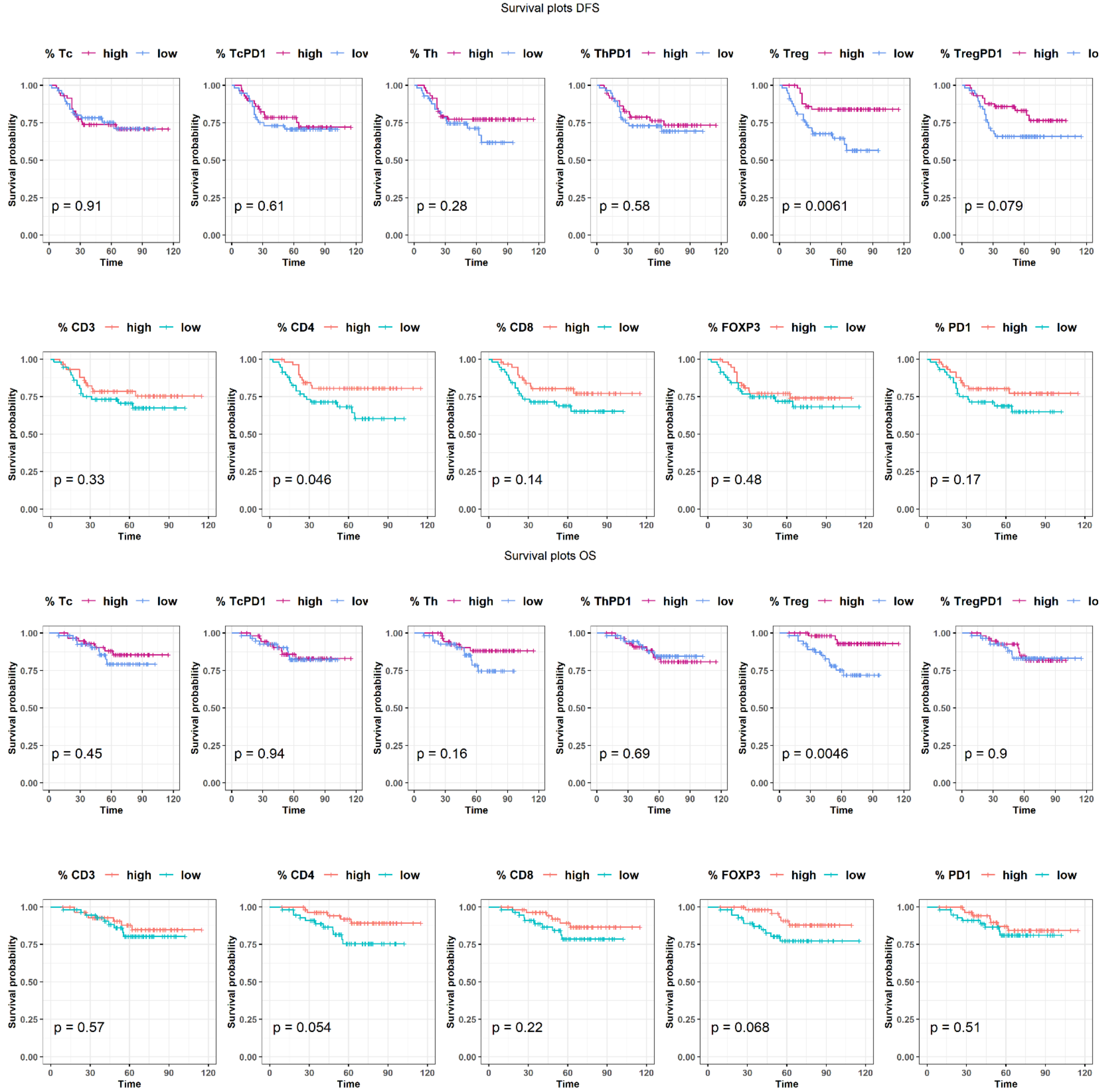

Supplementary Figure 8

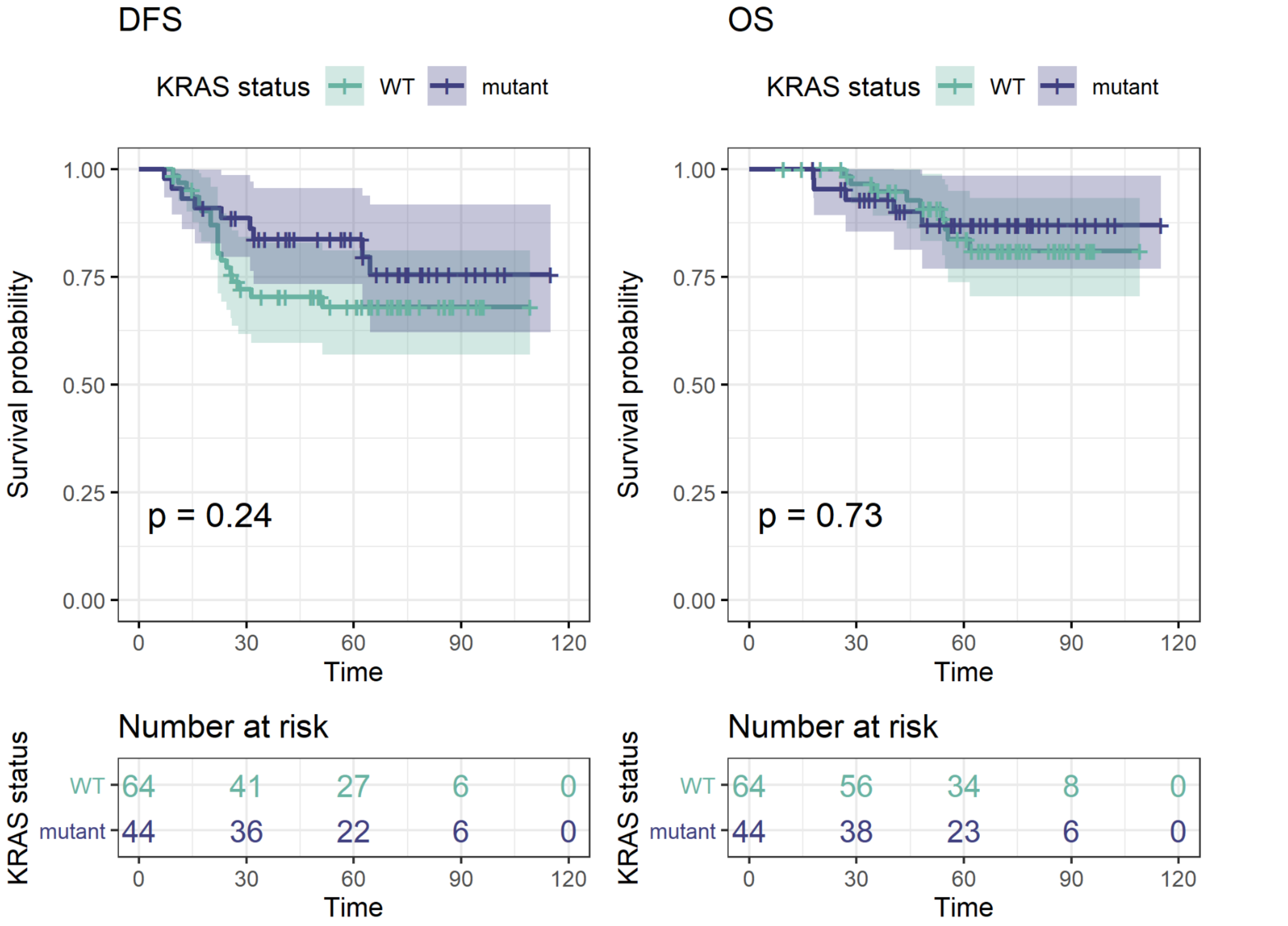

Supplementary Table 1: Antibodies, clones and conjugates used in this study.

| Target | Clone | Vendor | Catalog | Concentration (µg/mL) | Staining round |
|---|---|---|---|---|---|
| APAF-1 | 2E12 | Millipore | MAB3053 | 5 | 1 |
| Bak | D4E4 | Cell Signaling | 12105 | 5 | 5 |
| Bax | E63 | Abcam | ab216985 | 10 | 4 |
| BCL-2 | 124 | Lifespan | LS-C389442 | 5 | 1 |
| Bcl-xL | 7D9 | Thermo | MS-1334 | 10 | 6 |
| CA9 | polyclonal | Thermo | PA1-16592 | 15 | 11 |
| Caspase-3 (Pro+cleaved) | D3R6Y | Cell Signaling | 14214 | 10 | 3 |
| Caspase-9 | 96.1.23 | Santa Cruz | sc-56076 A647 | 5 | 2 |
| **CD3** | F7.2.38 | Dako | M7254 | 10 | 12 |
| **CD4** | EPR6855 | Abcam | ab181724 | 5 | 9 |
| **CD8** | C8/144B | Dako | M7103 | 5 | 7 |
| **CD45** | 2B11 + PD7/26 | Dako | M0701 | 10 | 10 |
| **Cytokeratin AE1** | AE1 | eBioscience | 14-9001 | 2.5 | 8 |
| **Cytokeratin PCK26** | PCK26 | Sigma | C1801 | 5 | 7 |
| **FOXP3** | 206D | Biolegend | 320014 | 10 | 8 |
| Glut-1 | EPR3915 | Abcam | ab196357 | 5 | 11 |
| HLA I | EMR8 5 | Abcam | ab70328 | 10 | 10 |
| Ki67 | SP6 | Zeta | Z2031 | 10 | 9 |
| MCL-1 | Y37 | Abcam | ab186822 | 10 | 2 |
| **NAKATPase** | EP1845Y | Abcam | ab167390 | 5 | 6 |
| **S6** | C-8 | Santa Cruz | sc-74459 A647 | 5 | 13 |
| Smac | 79-1-83 | Cell Signaling | 2954 | 10 | 4 |
| **PD1** | EPR4877(2) | Abcam | ab201825 | 5 | 12 |
| XIAP (API3) | polyclonal | Thermo | APH937 | 7.5 | 5 |

Supplementary Table 2: Hazard Ratios for single-marker and multi-marker T cell subsets for average of cores.

**Multi-Marker**

| % of Total | DFS | | | OS | | |
|---|---|---|---|---|---|---|
| **Stroma** | **HR** | **95% CI for HR** | **p.value** | **HR** | **95% CI for HR** | **p.value** |
| **Tc** | 0.62 | 0.3-1.3 | 0.2 | 0.62 | 0.22-1.7 | 0.36 |
| **TcPD1** | 0.4 | 0.11-1.4 | 0.15 | 0.92 | 0.29-3 | 0.89 |
| **Th** | 0.13 | 0.0055-3 | 0.2 | 0.1 | 0.0011-9.8 | 0.33 |
| **ThPD1** | 0.51 | 0.18-1.4 | 0.19 | 0.65 | 0.2-2.2 | 0.48 |
| **Treg** | 0.38 | 0.14-1 | 0.052 | 0.17 | 0.025-1.1 | 0.067 |
| **TregPD1** | 0.3 | 0.05-1.8 | 0.19 | 0.49 | 0.055-4.3 | 0.52 |
| **Epithelia-associated** | | | | | | |
| **Tc** | 0.097 | 0.0034-2.8 | 0.17 | 0.024 | 7.6e-05-7.5 | 0.2 |
| **TcPD1** | 0.066 | 0.0035-1.2 | 0.07 | 0.19 | 0.007-5.2 | 0.33 |
| **Th** | 3.90E+07 | 0.0026-6e+17 | 0.14 | 7.40E-12 | 3e-50-1.8e+27 | 0.57 |
| **ThPD1** | 0.27 | 0.011-6.5 | 0.42 | 0.1 | 0.00055-18 | 0.39 |
| **Treg** | 2.10E-15 | 1.1e-30-3.9 | 0.06 | 4.30E-18 | 2.7e-41-680000 | 0.14 |
| **TregPD1** | 3.50E-06 | 3.5e-15-3600 | 0.24 | 5.90E-15 | 2.7e-33-13000 | 0.13 |

**Single-Marker**

| % of Total | DFS | | | OS | | |
|---|---|---|---|---|---|---|
| **Stroma** | **HR** | **95% CI for HR** | **p.value** | **HR** | **95% CI for HR** | **p.value** |
| **CD3** | 0.87 | 0.71-1.1 | 0.16 | 0.94 | 0.75-1.2 | 0.56 |
| **CD4** | 0.87 | 0.74-1 | 0.1 | 0.86 | 0.69-1.1 | 0.21 |
| **CD8** | 0.76 | 0.57-1 | 0.07 | 0.83 | 0.58-1.2 | 0.3 |
| **FOXP3** | 0.78 | 0.56-1.1 | 0.14 | 0.67 | 0.39-1.2 | 0.15 |
| **PD1** | 0.5 | 0.2-1.2 | 0.12 | 0.83 | 0.33-2.1 | 0.7 |
| **Epithelia-associated** | | | | | | |
| **CD3** | 0.64 | 0.29-1.4 | 0.27 | 0.75 | 0.28-2 | 0.56 |
| **CD4** | 0.52 | 0.082-3.3 | 0.49 | 0.027 | 0.00015-4.8 | 0.17 |
| **CD8** | 0.34 | 0.1-1.1 | 0.081 | 0.36 | 0.071-1.8 | 0.21 |
| **FOXP3** | 0.00018 | 2.4e-09-13 | 0.13 | 6.10E-10 | 3.7e-20-10 | 0.077 |
| **PD1** | 0.052 | 0.0023-1.2 | 0.064 | 0.13 | 0.0035-4.9 | 0.27 |

Supplementary Table 3: Hazard Ratios for single-marker and multi-marker T cell subsets from immune hot-spot

| Multi-Marker | | | | | | |
|---|---|---|---|---|---|---|
| **% of Total** | **DFS** | | | **OS** | | |
| **Stroma** | **HR** | **95% CI for HR** | **p.value** | **HR** | **95% CI for HR** | **p.value** |
| **Tc** | 0.84 | 0.58-1.2 | 0.37 | 0.83 | 0.47-1.4 | 0.51 |
| **TcPD1** | 0.67 | 0.33-1.4 | 0.27 | 1 | 0.49-2.1 | 0.95 |
| **Th** | 0.59 | 0.17-2 | 0.4 | 0.31 | 0.026-3.6 | 0.35 |
| **ThPD1** | 0.79 | 0.46-1.4 | 0.39 | 0.8 | 0.4-1.6 | 0.54 |
| **Treg** | 0.52 | 0.27-0.98 | 0.042* | 0.25 | 0.061-1 | 0.052 |
| **TregPD1** | 0.59 | 0.24-1.5 | 0.25 | 0.8 | 0.28-2.3 | 0.68 |
| **Epithelia-associated** | | | | | | |
| **Tc** | 0.39 | 0.059-2.6 | 0.33 | 0.22 | 0.0099-4.9 | 0.34 |
| **TcPD1** | 0.32 | 0.063-1.6 | 0.16 | 0.33 | 0.035-3.1 | 0.33 |
| **Th** | 3300 | 5.2e-06-2.1e+12 | 0.43 | 0.052 | 4.2e-19-6.5e+15 | 0.88 |
| **ThPD1** | 0.9 | 0.27-3 | 0.86 | 0.5 | 0.05-4.9 | 0.55 |
| **Treg** | 6.90E-08 | 2e-17-240 | 0.14 | 1.70E-13 | 7.3e-32-410000 | 0.17 |
| **TregPD1** | 0.71 | 1.3e-05-40000 | 0.95 | 7.50E-11 | 2.5e-24-2300 | 0.14 |

| Single-Marker | | | | | | |
|---|---|---|---|---|---|---|
| **% of Total** | **DFS** | | | **OS** | | |
| **Stroma** | **HR** | **95% CI for HR** | **p.value** | **HR** | **95% CI for HR** | **p.value** |
| **CD3** | 0.91 | 0.8-1 | 0.17 | 0.94 | 0.8-1.1 | 0.45 |
| **CD4** | 0.92 | 0.83-1 | 0.11 | 0.89 | 0.76-1 | 0.13 |
| **CD8** | 0.86 | 0.71-1 | 0.12 | 0.83 | 0.62-1.1 | 0.2 |
| **FOXP3** | 0.77 | 0.59-1 | 0.052 | 0.66 | 0.43-1 | 0.066 |
| **PD1** | 0.7 | 0.41-1.2 | 0.21 | 0.88 | 0.47-1.6 | 0.68 |
| **Epithelia-associated** | | | | | | |
| **CD3** | 1 | 0.73-1.5 | 0.87 | 1.1 | 0.74-1.7 | 0.59 |
| **CD4** | 0.91 | 0.44-1.9 | 0.81 | 0.38 | 0.044-3.4 | 0.39 |
| **CD8** | 0.45 | 0.19-1.1 | 0.085 | 0.3 | 0.064-1.4 | 0.13 |
| **FOXP3** | 0.11 | 0.00019-61 | 0.49 | 7.90E-05 | 7.6e-11-81 | 0.18 |
| **PD1** | 0.2 | 0.029-1.3 | 0.097 | 0.23 | 0.018-2.8 | 0.25 |
